# Assessing the protective potential of H1N1 influenza virus hemagglutinin head and stalk antibodies in humans

**DOI:** 10.1101/478222

**Authors:** Shannon R. Christensen, Sushila A. Toulmin, Trevor Griesman, Lois E. Lamerato, Joshua G. Petrie, Emily T. Martin, Arnold S. Monto, Scott E. Hensley

**Author notes:** corresponding author: 402 Johnson Pavilion, 3610 Hamilton Walk, Philadelphia, PA 19104.

## Abstract

Seasonal influenza viruses are a major cause of human disease worldwide. Most neutralizing antibodies (Abs) elicited by influenza viruses target the head domain of the hemagglutinin (HA) protein. Anti-HA head Abs can be highly potent, but they have limited breadth since the HA head is variable. There is great interest in developing new universal immunization strategies that elicit broadly neutralizing Abs against conserved regions of HA, such as the stalk domain. Although HA stalk Abs can provide protection in animal models, it is unknown if they are present at sufficient levels in humans to provide protection against naturally-acquired influenza virus infections. Here, we quantified H1N1 HA head and stalk-specific Abs in 179 adults hospitalized during the 2015-2016 influenza virus season. We found that HA head Abs, as measured by hemagglutinin-inhibition (HAI) assays, were associated with protection against naturally-acquired H1N1 infection. HA stalk-specific serum total IgG titers were also associated with protection, but this association was slightly attenuated and not statistically significant after adjustment for HA head-specific Ab titers. We found higher titers of HA stalk-specific IgG1 and IgA Abs in sera from uninfected participants than from infected participants; however, we found no difference in sera *in vitro* antibody dependent cellular cytotoxicity activity. In passive transfer experiments, sera from participants with high HAI activity efficiently protected mice, while sera with low HAI activity protected mice to a lower extent. Our data suggest that human HA head and stalk Abs both contribute to protection against H1N1 infection.

**Importance:** Abs targeting the HA head of influenza viruses are often associated with protection from influenza virus infections. These Abs typically have limited breadth since mutations frequently arise in HA head epitopes. New vaccines targeting the more conserved HA stalk domain are being developed. Abs that target the HA stalk are protective in animal models, but it is unknown if these Abs exist at protective levels in humans. Here, we found that Abs against both the HA head and HA stalk were associated with protection from naturally-acquired human influenza virus infections during the 2015-2016 influenza season.

## Introduction

Seasonal influenza viruses cause annual epidemics worldwide. Although seasonal influenza vaccines usually provide moderate protection against circulating strains, vaccine effectiveness can be low when there are antigenic mismatches between vaccine strains and circulating strains (1, 2). Additionally, rare yet unpredictable influenza pandemics occur when novel influenza strains cross the species barrier and transmit in the human population (3).

Antibody (Ab)-mediated immunity is important for protecting against influenza virus infections (4). The viral membrane protein, hemagglutinin (HA), is the target for most anti-influenza virus neutralizing Abs (5–9). Most neutralizing HA Abs target the HA globular head domain and block virus attachment to sialic acid, the cellular receptor for influenza viruses. However, since the HA head is highly variable, HA head Abs generally exhibit poor cross-reactivity against antigenically drifted viral strains (10). Unlike the head domain, the stalk domain of HA is highly conserved between different influenza virus strains. Abs that target the HA stalk domain can prevent viral replication by inhibiting the pH-induced conformational changes of HA that are required for viral entry into the cell (11). Many HA stalk-specific Abs also protect by blocking HA maturation (11), inhibiting viral egress (12), or by mediating Ab dependent cellular cytotoxicity (ADCC) (13). Although HA stalk Abs are typically subdominant and are not thought to be as efficient as HA head Abs, HA stalk Abs can inhibit diverse influenza strains *in vitro* (14–17).

Conventional influenza vaccines effectively elicit HA head-reactive Abs, but not HA stalk Abs (18). As a result, influenza vaccine effectiveness is dependent on the similarity of the HA head of circulating influenza virus strains and the HA head of vaccine strains (19). Antigenic mismatch between influenza vaccine strains and circulating viral strains have been especially problematic during recent years (20, 21). To circumvent the potential for antigenic mismatch, as well as to prepare against new pandemic viral strains, there is great interest in developing new universal immunization strategies that would elicit broadly reactive Abs against conserved regions of HA, such as the stalk domain (22).

HA stalk Abs protect animals from group 1 and group 2 influenza A virus infections (14, 16, 23-29). For example, human anti-HA stalk monoclonal Abs (mAbs) protect mice from lethal pH1N1 infection following prophylactic or therapeutic passive transfers (23, 28), as well as against H5N1 (16, 24, 28) or H7N9 lethal dose challenge (27). Both the prophylactic passive transfer of a human anti-HA stalk mAb or the elicitation of HA stalk-specific Abs by chimeric HA vaccination decreased viral loads in ferrets following pH1N1 infection (25). Additionally, passive transfer of human sera from H5N1 vaccinees protects mice from lethal pH1N1 infection (26), and this protection is likely mediated by HA stalk Abs. Passive transfer of broadly neutralizing HA stalk-specific mAbs against group 2 influenza A viruses also protects mice against heterosubtypic H3 viruses (29) and heterologous H3 and H7 viruses (14). Vaccine strategies designed to elicit HA stalk Abs in humans are currently being pursued (30-32). These strategies include sequential immunizations with chimeric HAs (19, 33), immunization with ‘headless’ HA antigens (30, 34, 35), and immunizations with mRNA-based vaccines expressing HA (32).

Despite the recent interest in developing new HA stalk-based vaccines, the amount of HA stalk Abs required to protect humans from influenza virus infections and influenza-related disease has not been established. A recent human pH1N1 challenge study demonstrated that HA stalk Ab titers are associated with reduced viral shedding, but are not independently associated with protection against influenza infection (36). While human influenza virus challenge studies are valuable, they have some limitations. For example, high doses of virus are used in these studies (37, 38), large numbers of individuals are typically pre-screened for certain immunological attributes prior to entering these studies (39), and the pathogenesis of infection differs from that of a natural infection, including key sites of viral replication (38, 40). Serological studies of individuals who naturally acquire influenza virus infections can also be used to identify specific types of Abs that are associated with protection. Here, we present a serological study to determine if serum HA head and stalk Abs are associated with protection against naturally-acquired H1N1 infection.

## Results

### HA head and stalk Abs are associated with protection against H1N1 infection

We analyzed sera collected from 179 participants enrolled in a hospital-based study during the 2015-16 influenza season (Supplemental Table 1). Adults hospitalized at the University of Michigan Hospital (Ann Arbor, MI, USA) or Henry Ford Hospital (Detroit, MI, USA) were enrolled according to a case definition of within ≤10 days of acute respiratory illness onset and subsequently tested for influenza by RT-PCR. Serum specimens collected at hospital admission were obtained for estimation of pre/early-infection antibodies; 58% of specimens included in this analysis were collected within 3 days of illness onset (41). We analyzed serum samples from 62 hospitalized individuals that had PCR-confirmed H1N1 influenza virus infections. Serum samples from 117 controls were selected from hospitalized individuals that had other respiratory diseases not caused by an influenza virus infection matching on age category (18 – 49 years, 50 – 64 years, ≥65 years) and influenza vaccination status.

**Table 1.**
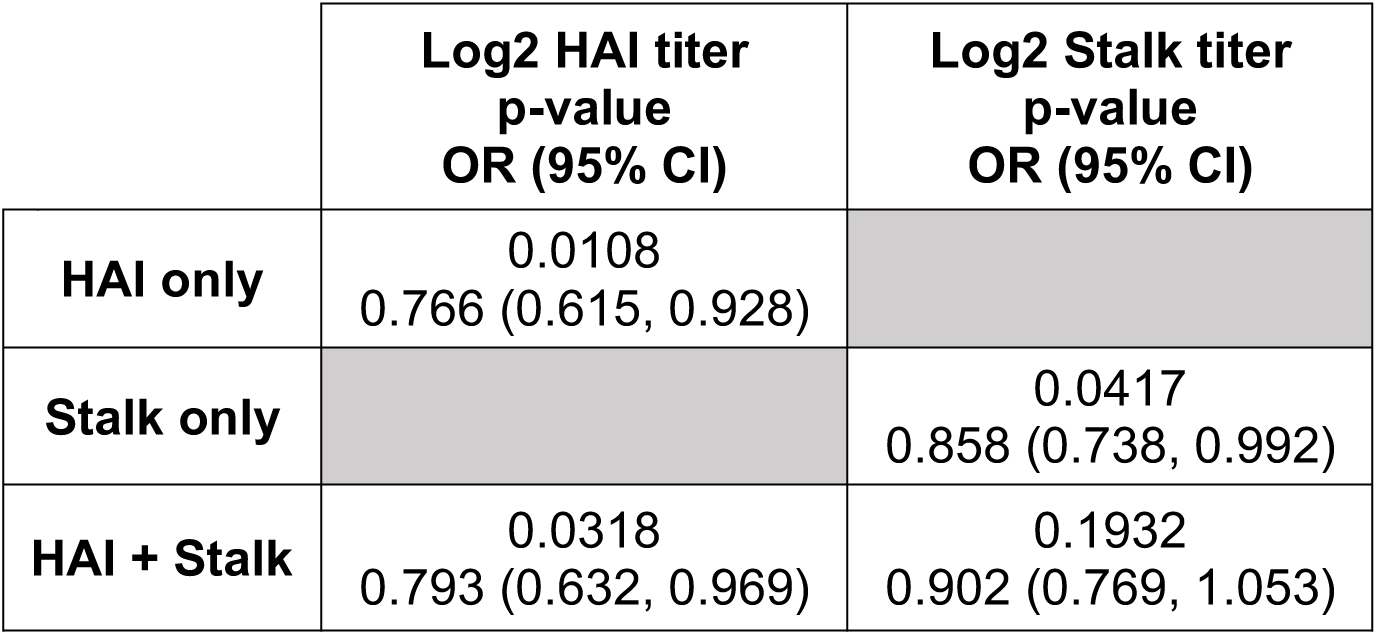
Logistic regression modeling of HA head and stalk antibodies association with protection. Logistic regression analyses using both unadjusted (HAI only and Stalk only) and adjusted (HAI + Stalk) models. Values represent log2 geometric mean titers of two independent experiments.

|  | **Log2 HAI titer**<br>**p-value**<br>**OR (95% CI)** | **Log2 Stalk titer**<br>**p-value**<br>**OR (95% CI)** |
| --- | --- | --- |
| **HAI only** | 0.0108<br>0.766 (0.615, 0.928) |  |
| **Stalk only** |  | 0.0417<br>0.858 (0.738, 0.992) |
| **HAI + Stalk** | 0.0318<br>0.793 (0.632, 0.969) | 0.1932<br>0.902 (0.769, 1.053) |

We quantified relative serum titers of HA head-specific Abs against the predominant 2015-2016 H1N1 strain using hemagglutination inhibition (HAI) assays (Fig. 1A). HAI assays detect HA head-specific Abs that prevent influenza virus mediated cross-linking of red blood cells (42, 43). We found that HAI titers were associated with protection against H1N1 infection in logistic regression models (Table 1). We observed a 23.4% reduction in H1N1 infection risk with every 2-fold increase in HAI titer. Previous studies reported that a 1:40 HAI titer is associated with 50% protection from experimental human influenza infections (44). Consistent with this, over 21% of non-H1N1 infected cases possessed >40 HAI titer, while only ~3% of H1N1 infected cases possessed >40 HAI titer (Supplemental Figure 1).

**Figure 1.**
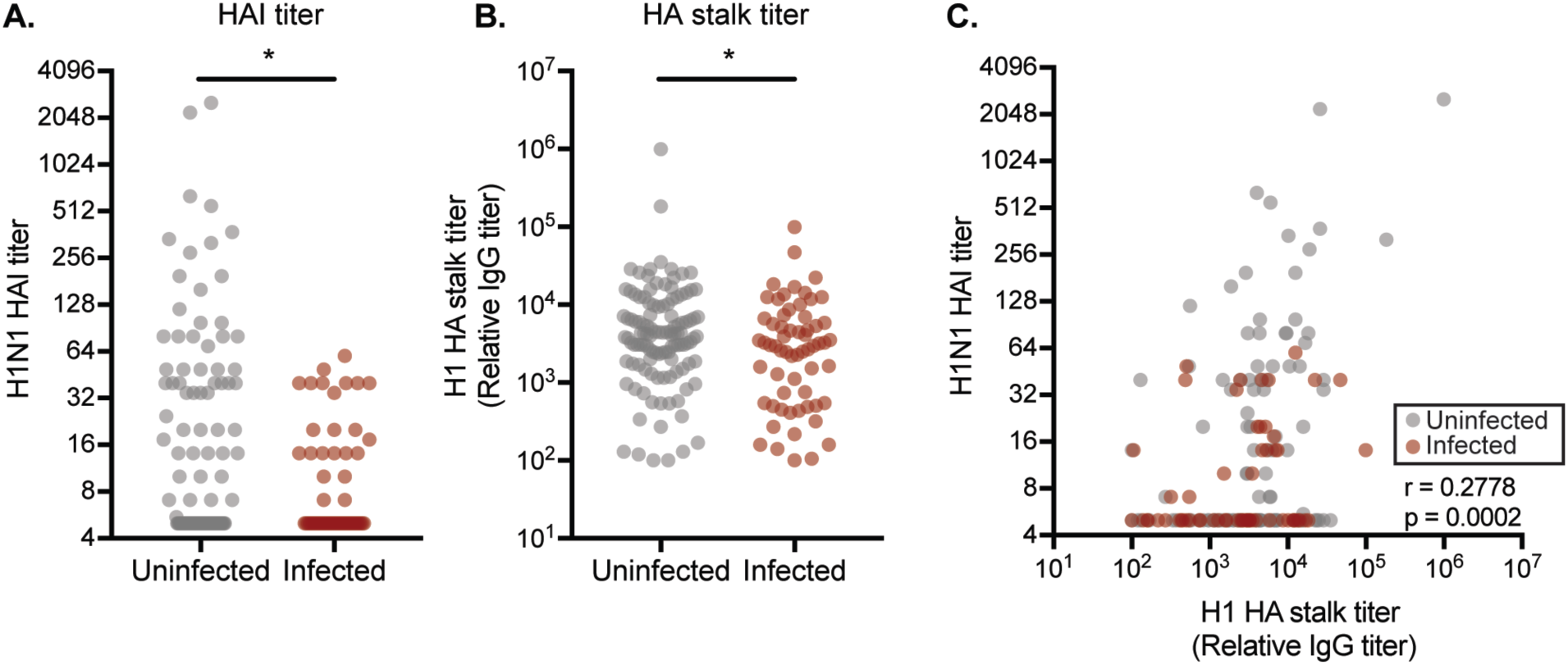
HA head and stalk antibodies are associated with protection. ***1A:*** HAI assays were completed using sera from uninfected (grey) and infected (red) individuals. HAI titers are associated with protection against H1N1 infection (p = 0.0108, logistic regression of log2 geometric mean titers of two independent experiments).***1B:*** ELISA assays using ‘headless’ HA constructs were completed using sera from uninfected (grey) and infected (red) individuals. HA stalk-specific Abs are associated with protection against influenza infection (p = 0.0417, logistic regression analysis using log2 geometric mean titers of two independent experiments.) ***1C.*** HA head Abs measured by HAI and HA stalk titers measured by ELISA using ‘headless’ HA stalk constructs are weakly, though significantly, correlated (r = 0.2778, p = 0.0002, Spearman Correlation using log2 geometric mean titers of two independent experiments for each measurement). In all figure panels each circle represents a serum sample from a single individual.

Next, we quantified relative titers of H1 stalk IgG Abs using ELISAs coated with ‘headless’ H1 proteins (Fig 1B). Similar to HAI titers, we found that H1 stalk titers were associated with protection against H1N1 infection in logistic regression models (Table 1). We observed a 14.2% reduction in H1N1 infection risk with every 2-fold increase in H1 stalk titer. Whereas HAI titers >40 were sharply associated with protection (Fig 1A, C), there was no clear HA stalk IgG titer cutoff that was associated with protection in our study (Fig. 1B, C). Samples with the highest HA stalk IgG titers were interspersed among the ‘uninfected’ and ‘infected’ groups (Fig. 1B, C).

Although both HAI titers and HA stalk IgG titers were associated with H1N1 protection in unadjusted models (Table 1), only HAI titers remained statistically associated with protection in adjusted models (Table 1). HA stalk IgG-associated protection lost significance when adjusting for HAI titers; however, the overall reduction in odds of infection for each 2-fold increase in titer remained roughly the same between the unadjusted and adjusted models for both HAI (23.4% to 20.7) and stalk Ab titers (14.2% to 9.8%), respectively (Table 1).

We completed several experiments to validate our HA stalk IgG data. ‘Headless’ H1 proteins are engineered to possess only the HA stalk domain and not the HA globular head domain (30). We completed experiments with monoclonal Abs (mAbs) to verify that HA stalk-reactive Abs bind to ‘headless’ H1 proteins in ELISAs. We found that the H1 head-specific EM-4C04 mAb bound efficiently to a full-length H1 HA protein, but failed to bind to our ‘headless’ H1 protein, while the H1 stalk-specific 70-1F02 mAb bound to each construct similarly (Figure 2A-B). We used two additional methods to verify that ‘headless’ HA-based ELISAs accurately quantify HA stalk-reactive Abs. First, we measured Ab binding to a full-length HA chimeric protein that possessed an “exotic” head domain from an H6 virus fused to the H1 stalk (abbreviated c6/H1). Since H6 viruses have never circulated in the human population, most human Abs that bind to this recombinant HA target the HA stalk domain (19). We found similar relative HA stalk Ab levels when we used the c6/H1 HA-based ELISAs compared to ‘headless’ HA-based ELISAs (Fig. 2C). We also quantified HA stalk Ab levels using a competition ELISA. For these experiments we determined the amount of serum Abs that were required to prevent the binding of a biotinylated HA stalk-specific mAb (70-1F02). The 70-1F02 mAb recognizes a conformationally dependent epitope that spans the HA1 and HA2 subunits (15, 17, 28, 45, 46). We found that 70-1F02-based competition assay titers correlated strongly with the ‘headless’ HA-based ELISA titers (Fig. 2D).

**Figure 2.**
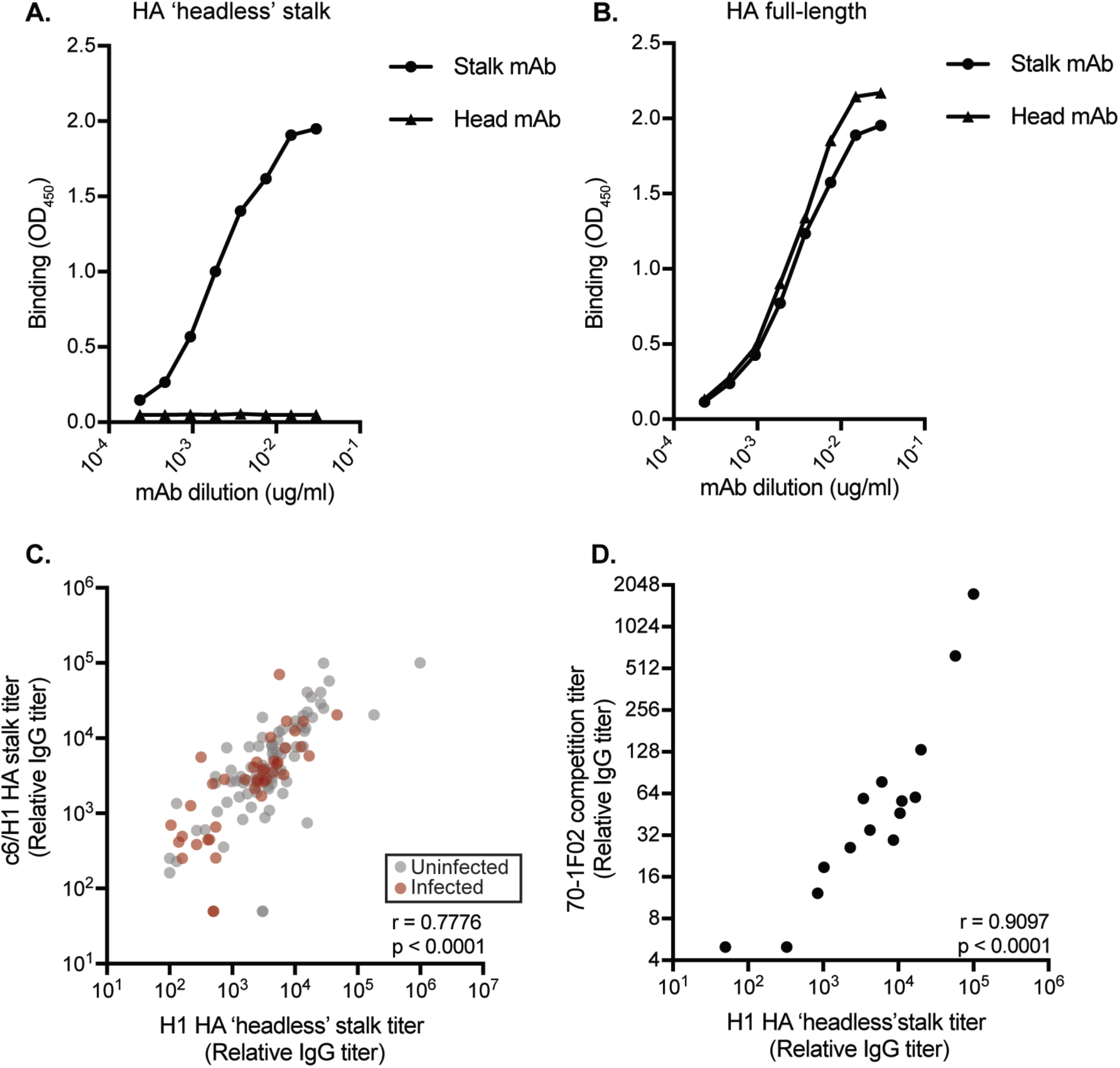
Validation of ‘headless’ H1 HA stalk construct. We completed additional ELISAs using the 70-1F02 HA stalk mAb and the EM-4C04 HA head Ab and plates coated with ‘headless’ HA (**2A**) or full length HA (**2B**). Graphs depict representative results from two independent experiments. ***2C:*** We quantified HA stalk Abs using ELISA plates coated with c6/H1 proteins. HA stalk titers measured by ELISA using c6/H1 or ‘headless’ HA stalk constructs were tightly correlated (r = 0.7776, p < 0.0001, Spearman Correlation using log2 geometric mean titers of two independent experiments). ***2D:*** We completed competition assays using the conformationally-dependent 70-1F02 mAb. 70-1F02 competition titers are tightly correlated with overall HA stalk Ab titers in both infected and uninfected individuals (r = 0.9097, p < 0.0001, Spearman Correlation using log2 geometric mean titers of two independent experiments). In **2C-2D** each circle represents a serum sample from a single individual.

### HA stalk IgG1 and IgA Abs are associated with protection

Some HA stalk Abs mediate protection through non-neutralizing mechanisms that involve processes such as ADCC (47). IgG1 and IgG3 Ab subtypes are efficient at inducing ADCC, whereas IgG2 and IgG4 are not (48). We quantified relative IgG1, IgG2, and IgG3 HA stalk Abs in serum from a subset of participants using ELISAs coated with ‘headless’ H1 proteins. In all participants, the majority of HA stalk IgGs were IgG1 (Fig. 3A), consistent with previous reports (49, 50). Total HA stalk IgG titers closely correlated with HA stalk IgG1 titers (Fig. 3B). Similar to total HA stalk IgG titers (Fig. 1B), we found that H1 stalk IgG1 titers were associated with H1N1 protection in logistic regression models (Fig. 3A & Table 2). Very low levels of IgG2 and IgG3 HA stalk Abs were detected in serum (Fig. 3A). We did not measure levels of IgG4 HA stalk Abs since IgG4 is not prevalent among anti-influenza virus human Abs (49). It is important to note that titers of each isotype are directly comparable in our experiments since we used control mAbs in each ELISA (based on the CR9114 HA stalk mAb (51, 52)) that were engineered to possess the same variable regions coupled to different constant regions (Supplemental Figure 2).

**Table 2.** Logistic regression modeling of HA stalk antibody isotypes association with protection. Logistic regression analyses were performed using unadjusted models. Values represent log2 geometric mean titers of two independent experiments.

|  | **Log2 Stalk IgG1**<br>**titer p-value**<br>**OR (95% CI)** | **Log2 Stalk IgA titer**<br>**p-value**<br>**OR (95% CI)** |
| --- | --- | --- |
| **Unadjusted** | 0.0124<br>0.801 (0.669, 0.950) | 0.0433<br>0.846 (0.716, 0.991) |

**Figure 3.**
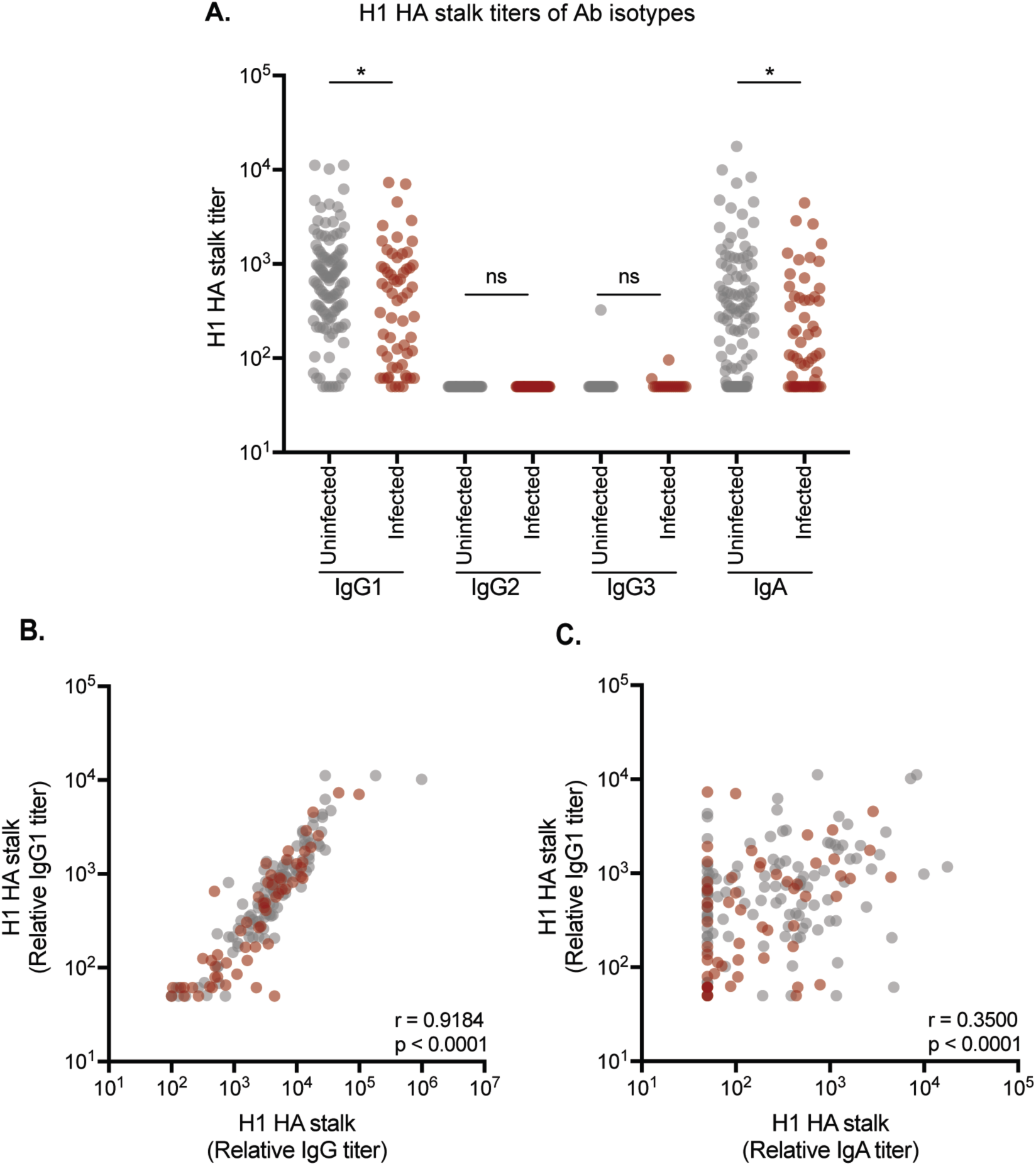
HA stalk-specific serum IgG1 and IgA are associated with protection. ***3A:*** ELISAs were completed to quantify the levels of IgG1, IgG2, IgG3, and IgA HA stalk Abs in each serum sample. HA stalk-specific IgG1 and IgA are associated with protection against influenza infection (p = 0.0433 and p = 0.0124, respectively. Logistic regression analysis using log2 geometric mean titers of two independent experiments.) ***3B.*** IgG1 HA stalk Ab titers closely correlated with total IgG HA stalk Ab titers (r = 0.9184, p < 0.0001, Spearman Correlation using log2 geometric mean titers of two independent experiments). ***3C.*** IgA HA stalk Ab titers moderately correlated with IgG1 HA stalk Ab titers (r = 0.3500, p < 0.0001, Spearman Correlation using log2 geometric lmean titers of two independent experiments). In all figure panels each circle represents a serum sample from a single individual.

We next evaluated serum HA stalk IgA Abs, since IgA Abs can be important for controlling respiratory infections. For example, mucosal IgA potently reduces the risk of influenza transmission events in guinea pigs in a dose-dependent manner (53) and suppresses the extracellular release of virus from infected cells (54). Further, anti-HA stalk mAbs engineered on an IgA backbone neutralize virus more effectively compared to when they are engineered on an IgG backbone (55). We did not have access to respiratory secretions, but we did measure levels of HA stalk monomeric IgA in the serum from a subset of participants. Similar to HA stalk total IgG (Fig. 1B) and IgG1 (Fig. 3A) titers, we found that serum HA stalk IgA titers were associated with H1N1 protection in logistic regression models (Fig. 3A & Table 2). IgG1 and IgA titers were moderately, though significantly, correlated (Fig. 3C).

### Functionality of Abs from infected and uninfected individuals

Abs against the HA head and HA stalk can neutralize or limit virus replication through distinct mechanisms (11-14, 45-47, 56, 57). For example, most Abs that target epitopes near the receptor binding domain of the HA head block virus binding and neutralize virus *in vitro* and *in vivo* (56). Some HA stalk Abs can directly neutralize virus, but the majority of HA stalk Abs require Fc receptor engagement for protection *in vivo* (15, 47, 58). Neutralizing HA stalk Abs typically inhibit HA conformational changes required to mediate fusion of the virus and cellular membranes (11, 14, 45, 46). Other HA stalk Abs can prevent subsequent viral expansion at later stages of infection by inhibiting HA maturation (11) and viral egress (12).

We completed experiments to assess the *in vitro* and *in vivo* protective potential of serum Abs from infected and uninfected individuals. First, we performed *in vitro* neutralization assays using GFP-reporter influenza viruses (59). We generated H1N1 viruses possessing genes encoding enhanced green fluorescent protein (eGFP) in place of most of the PB1 gene segment. The eGFP segment retained the noncoding and 80 terminal coding nucleotides, allowing this segment to be efficiently and stably packaged into virions. Neutralization assays were completed with these viruses in cell lines that stably expressed PB1. We detected *in vitro* neutralization titers in serum from approximately half of the participants that we tested. We found that *in vitro* neutralization titers were significantly associated with protection in logistic regression models (Fig. 4A). As expected, serum samples with the highest HAI titers had high *in vitro* neutralization titers (Fig. 4B), whereas serum samples with the highest HA stalk titers had more variable *in vitro* neutralization titers (Fig. 4C).

**Figure 4.**
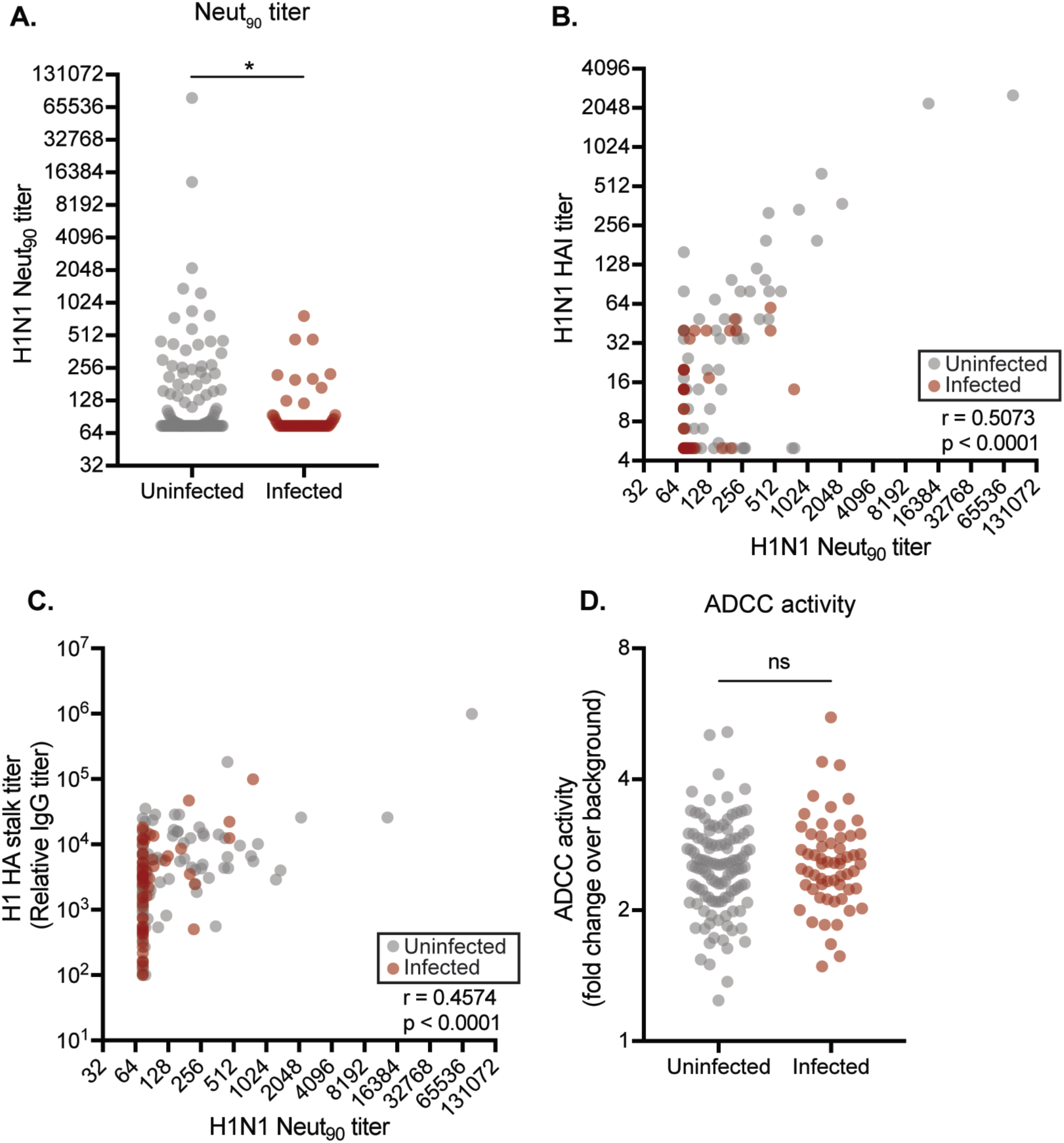
*In vitro* functionality of HA Abs from infected and uninfected individuals. ***4A:** In vitro* neutralization assays were completed with sera from uninfected and infected individuals. *In vitro* neutralization titers are associated with protection against influenza infection (p = 0.0185, logistic regression analysis using log2 geometric mean titers of two independent experiments). ***4B:*** HAI titers correlate strongly with neutralization titers (r = 0.5073, p < 0.0001, Spearman correlation using log2 geometric mean titers of two independent experiments). ***4C:*** HA stalk titers also correlate with neutralization titers (r = 0.4574, p < 0.0001, Spearman correlation using log2 geometric mean titers of two independent experiments). ***4D:*** ADCC assays were completed using sera from uninfected and infected individuals. ADCC activity is not associated with protection against influenza infection (p = 0.4160, logistic regression analysis using log2 geometric mean titers of three independent experiments).

Next, we completed *in vitro* ADCC assays using serum from a subset of participants. For these assays we incubated HA-expressing 293T cells with serum, and then added human peripheral blood mononuclear cells (PBMCs). We then measured CD107a (LAMP1) expression on CD3^-^CD56^+^ NK cells by flow cytometry. CD107a is a sensitive NK cell degranulation marker whose expression levels strongly correlate with cytokine production and cytotoxicity by NK cells in response to Ab-Fc receptor engagement (60). Unlike *in vitro* neutralization titers (Fig. 4C), ADCC activity was not associated with protection in logistic regression models (Fig. 4D).

Finally, we completed passive transfer experiments in mice. For these experiments we passively transferred human sera into mice that have been engineered to possess human Fc-receptors (61) so that we could accurately assess the protective effects mediated by human Fc-FcR interactions. We passively transferred pooled sera from uninfected individuals that had high (>40) HAI titers (abbreviated as uninfected-HAI^high^) and uninfected individuals that had low (≤ 40) HAI titers (abbreviated as uninfected-HAI^low^). We also passively transferred pooled sera from infected individuals, all of whom had low (≤ 40) HAI titers (abbreviated as infected-HAI^low^). For these experiments, equal volumes of human sera were transferred for each experimental condition. Mice were challenged with a sub-lethal dose of H1N1 four hours after sera transfer and body weights were monitored for 15 days (Figure 5A). Mice that received sera from uninfected-HAI^high^ participants were fully protected against H1N1 infection. Mice that received sera from HAI^low^ participants, whether from uninfected or infected individuals, were moderately protected against H1N1 infection, though these sera conferred significantly less protection when compared to sera from HAI^high^ participants (Figure 5B and Supplemental Table 2). Since there were different amounts of HA Abs in sera from uninfected-HAI^high^ participants, uninfected-HAI^low^ participants, and infected participants (Fig 5C), it is unclear if the differences in our passive transfer experiments were due to differences in overall HA Ab titers or differences in HA head and stalk Ab ratios. To address this, we repeated passive transfer experiments after adjusting sera amounts so that equal amounts of HA Abs were passively transferred in each experimental group. Similar to what we found in our initial passive transfer experiment, sera from uninfected-HAI^high^ participants protected mice better compared to sera from uninfected-HAI^low^ participants and infected participants (Fig. 5D and Supplemental Table 3). Interestingly, sera from uninfected-HAI^low^ participants protected mice better compared to sera from infected participants after adjusting sera amounts based on HA Ab titers (Fig. 5D and Supplemental Table 3). Taken together, these data suggest that human sera with high HAI activity efficiently protect *in vivo*, while human sera with low HAI activity also protect *in vivo*, albeit to a lower extent.

**Figure 5.**
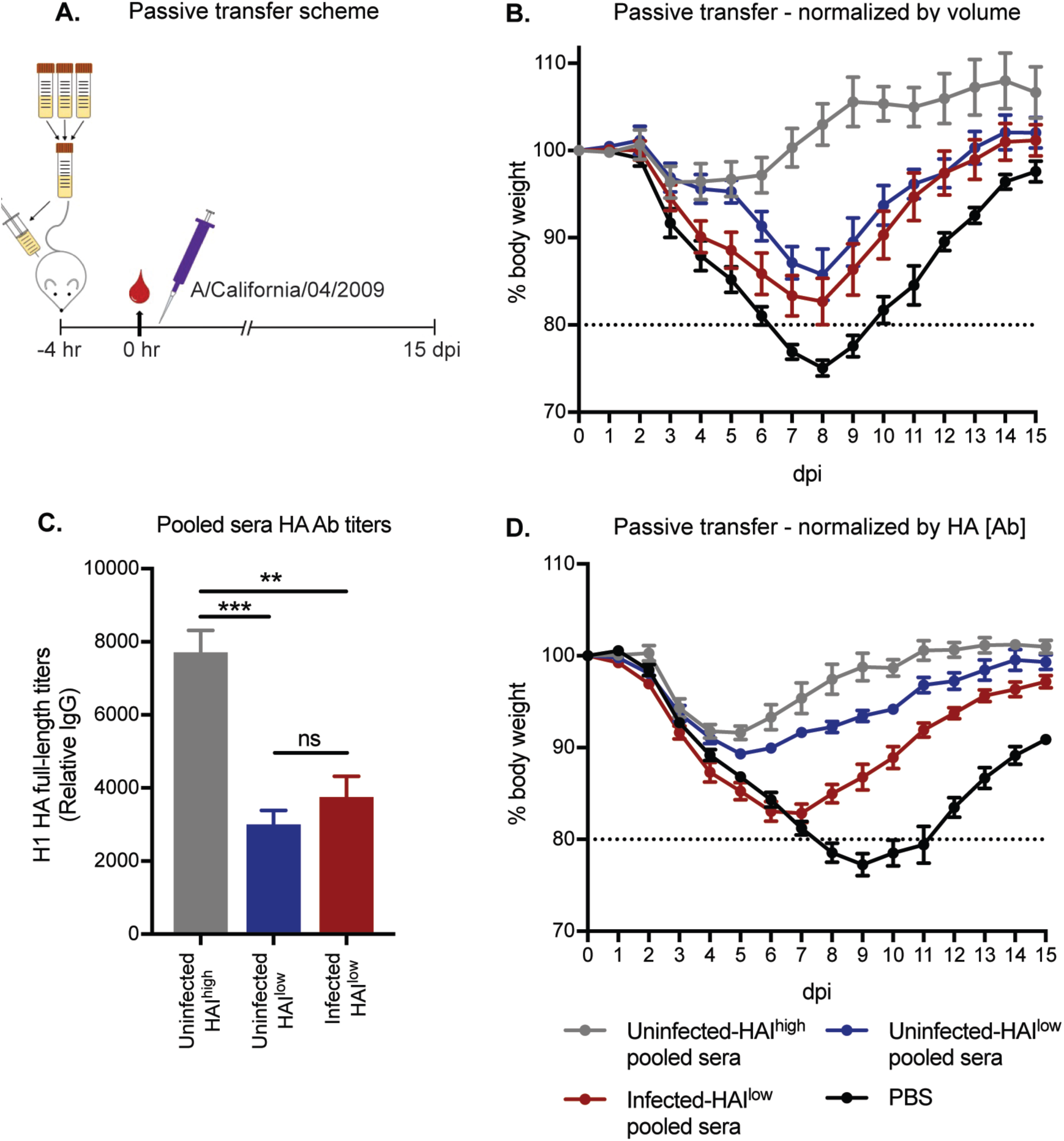
HA head and stalk antibodies confer protection from severe disease and mortality *in vivo*. ***5A:*** Passive transfer experiment design and timeline. Sera was stratified by HAI titer and infection status, pooled, and transferred I.P. to humanized Fc-receptor mice four hours before challenge with A/California/04/2009. Weights were measured daily for 15 days. ***5B:*** We transferred equal volumes of sera into each mouse for our initial experiments. Mice that received uninfected-HAI^high^ sera were completed protected against infection (grey line). Mice that received HAI^low^ sera (uninfected or infected – blue and red lines, respectively) were protected against mortality, but were significantly less protected compared to the mice that received HAI^high^ sera (+/-SEM, 1-way ANOVA analysis performed for each dpi based on %-weight lost relative to starting weight using two independent experiments with 6 mice/group. Results listed in Supplemental Table 2). ***5C:*** We completed ELISAs to quantify total H1-reactive Abs in each of our pooled sera samples and found that there were different amounts of HA Abs in each sample. **5D:** We repeated passive transfer studies after normalizing sera amounts so that equal amounts of HA Abs were transferred. Mice that received uninfected-HAI^high^ or uninfected-HAI^low^ pooled sera (grey and blue lines, respectively) were protected similarly against severe influenza disease and mortality, with mice that received uninfected HAI^high^ pooled sera recovering more quickly than mice that received uninfected-HAI^low^ pooled sera (+/-SEM, 1-way ANOVA analysis performed for each dpi based on %-weight lost relative to starting weight using one independent experiment with 6 mice/group. Results listed in Supplemental Table 3).

## Discussion

Observational studies can be useful in identifying Ab types that are associated with protection from influenza virus infection. Here, we found that both HA head and stalk Abs appeared to be associated with preventing H1N1 hospitalizations during the 2015-2016 season. We found that the effect size of HAI-associated protection (23.4% reduced risk of infection for every 2-fold increase in titer) was larger than the effect size of HA stalk Ab-associated protection (14.2% reduced risk of infection for every 2-fold increase in titer). In our study, HAI titers were independently associated with protection in adjusted models, however, HA stalk Abs were not. However, the effects of both HAI and HA stalk Ab titers were only slightly attenuated in our adjusted model and it is possible that our relatively small sample size limited our ability to detect an independent association between HA stalk titers and protection.

There are several limitations to our study. Since our sample size was relatively small, we only evaluated the contribution of Abs to the HA head and stalk. Larger studies will be required to independently evaluate other immune correlates of protection. For example, it will be important for future studies to evaluate the relationship between HA head and stalk Ab-associated protection and neuraminidase (NA) Ab-associated protection. Recent studies have shown that NA Abs are associated with protection in an H1N1 challenge cohort (36), and NA Abs were also identified as an independent correlate of protection in a controlled vaccine efficacy study (62). It will be critical to determine if NA Ab-associated protection is independent of the protective effects of HA head and stalk Abs.

It should be noted that participants in our study were likely admitted to the hospital at varying days post-infection. While most blood specimens were collected relatively early (<=3 days after symptom onset), we cannot exclude that some participants in our studies were infected for a prolonged period of time before being admitted to the hospital. This raises the possibility that some participants may have already mounted *de novo* Ab responses to H1N1 infection, which could potentially convolute the analyses of Ab types associated with protection. While this is a possibility, it is less of a concern since we found that all infected individuals have very low HAI titers. If our infected participants were making *de novo* Abs responses, we would anticipate that some of them would have high HAI titers to the infecting H1N1 virus. In addition, Ab titers do not typically increase as days from symptom onset to blood specimen collection increases (41), which suggests that samples used in this study were collected prior to the generation of *de novo* Ab responses against the infecting virus.

It is interesting that *in vitro* neutralization titers (Fig. 4A), but not ADCC titers (Fig. 4D), were associated with H1N1 protection. *In vitro* neutralization activity is mainly driven by HA head Abs (5–9), whereas HA stalk Abs are more effective at ADCC (47). It should be noted that HA stalk IgG1 and IgA Abs have been shown to mediate phagocytosis with innate cellular partners (63), which could prove to be an important mechanism of protection by HA stalk Abs, and should be considered in future studies. HA head Abs were associated with greater protection in our cohorts compared to stalk Abs (Table 1) and these Abs conferred superior protection compared to HA stalk Abs when passively transferred into mice (Fig B and D). Interestingly, serum from HAI^low^ uninfected participants protected mice better compared to serum from HAI^low^ infected participants in passive transfer studies after normalizing total HA Ab amounts in each transfer condition. While these data suggest that HA stalk Abs can confer protection *in vivo*, we cannot rule out that other immune components (such as NA Abs) contributed to protection in these experiments.

Taken together, our findings provide important new insights into the prevalence and functionality of HA head and stalk Abs in humans. Future studies that tease out the interdependence of HA head and stalk Abs, as well as Abs and T cells against other viral antigens, will be useful in guiding the development of new universal influenza vaccine antigens.

## Materials and Methods

### Human Subjects

During the 2015-2016 influenza season, adult (≥18 years) patients hospitalized for treatment of acute respiratory illnesses at the University of Michigan Hospital in Ann Arbor, MI and Henry Ford Hospital in Detroit, MI were prospectively enrolled in a case-test negative design study of influenza vaccine effectiveness. All participants provided informed consent and were enrolled ≤10 days from illness onset during the period of influenza circulation (January-April 2015-2016). Participants completed an enrollment interview and had throat and nasal swab specimens collected and combined for influenza virus identification. Influenza vaccination status was defined by self-report and documentation in the electronic medical record and Michigan Care Improvement Registry (MCIR). When available, clinical serum specimens collected as early as possible after hospital admission were retrieved; all specimens were collected ≤10 days from illness onset based on the enrollment case definition. Studies involving human adults were approved by the Institutional Review Boards of University of Michigan and University of Pennsylvania. All experiments (HAI, ELISAs, *in vitro* neutralization assays, ADCC assays, and passive transfers) were completed at the University of Pennsylvania using pre-existing and de-identified sera.

### Viruses

Viruses possessing A/California/07/2009 HA and NA or A/HUP/04/2016 HA and NA were generated by reverse genetics using internal genes from A/Puerto Rico/08/1934. Viruses were engineered to possess the Q226R HA mutation, which facilitates viral growth in chicken eggs. Viruses were grown in fertilized chicken eggs and the HA gene was sequenced to verify that additional mutations did not arise during propagation. We isolated the A/HUP/04/2016 virus from respiratory secretions obtained from a patient at the Hospital of the University of Pennsylvania in 2016. For this process, de-identified clinical material from the Hospital of University of Pennsylvania Clinical Virology Laboratory was added to Madin-Darby canine kidney (MDCK) cells (originally obtained from the National Institutes of Health) in serum-free media with L-(tosylamido-2-phenyl) ethyl chloromethyl ketone (TPCK)-treated trypsin, HEPES and gentamicin. Virus was isolated form the MDCK-infected cells 3 days later. We extracted viral RNA and sequenced the HA gene of A/HUP/04/16.

### Recombinant HA Proteins

Plasmids encoding the recombinant ‘headless’ HA stalk were provided by Adrian McDermott and Barney Graham from the Vaccine Research Center at the National Institutes of Health. The ‘headless’ HA stalk protein was expressed in 293F cells and purified using Ni-NTA agarose (Qiagen, Mat# 1018244) in 5 ml polypropylene columns (Qiagen, Cat# 34964), washed with pH 8 buffer containing 50 mM Na2HCO3 + 300 mM NaCl + 20 mM imidazole, then eluted using pH 8 buffer containing 50 mM Na2HCO3 + 300 mM NaCl + 300 mM imidazole. Purified protein was buffer exchanged into PBS (Corning, Ref# 21-031-CM). Following purification, the ‘headless’ HA stalk proteins were biotinylated using the Avidity BirA-500 kit (Cat# BirA500) and stored in aliquots at −80C. Plasmids encoding the recombinant chimeric (c6/H1) HA were provided by Florian Krammer (Mt. Sinai). The detailed protocol for expression of this protein is published elsewhere (64). In brief, the c6/H1 HA protein was expressed in High Five baculovirus cells and purified using the same methods referenced for the ‘headless’ HA stalk protein. Purified protein was buffer exchanged into PBS (Corning, Ref# 21-031-CM) and stored in aliquots at −80C.

### mAbs

Plasmids encoding the human mAbs EM-4C04, 70-1F02, and CR9114 IgG1 isotypes were provided by Patrick Wilson at the University of Chicago. The heavy chain constant regions for IgG2, IgG3, and IgA (sequences listed below) were synthesized as a gBlock by IDT and cloned into the pSport6 vector containing the heavy chain of CR9114. All mAbs were expressed in 293T cells and purified four days post infection using NAb protein A/G spin kits (Thermo Fisher, Cat# 89950) for the IgG isotypes or using peptide M agarose (InvivoGen, Cat# gel-pdm-2) for the IgA isotype.

#### IgG2

CGCATGATGCGTCGACCAAGGGTCCTAGCGTTTTCCCGCTCGCACCTTGTAGTCGGAGCACCTCCGAATCTACGGCGGCGCTCGGATGTCTGGTTAAGGATTACTTTCCTGAACCTGTTACTGTATCTTGGAATTCAGGAGCACTGACATCTGGTGTACATACTTTTCCAGCGGTTTTGCAGTCATCTGGTCTTTATTCCCTGTCCAGTGTGGTAACAGTACCATCCTCAAACTTTGGAACTCAGACCTATACCTGCAATGTGGACCACAAGCCATCCAATACAAAAGTCGATAAGACTGTCGAGCGGAAGTGCTGTGTCGAATGCCCTCCCTGCCCCGCTCCGCCGGTTGCAGGGCCAAGTGTATTTCTTTTTCCACCAAAACCAAAAGATACGCTTATGATATCTCGCACGCCTGAAGTAACCTGCGTAGTCGTTGATGTAAGTCACGAGGATCCCGAAGTTCAATTCAATTGGTATGTAGATGGCGTTGAAGTGCATAATGCAAAGACCAAACCTAGAGAAGAACAATTCAATAGTACCTTTCGCGTGGTTAGCGTACTCACAGTCGTCCACCAGGATTGGCTGAATGGGAAGGAGTACAAATGCAAGGTCTCTAACAAAGGTCTTCCGGCCCCCATAGAAAAAACGATCAGTAAGACCAAGGGGCAGCCCAGAGAGCCACAGGTTTATACGTTGCCTCCGTCTCGCGAGGAAATGACTAAAAACCAGGTCAGCCTGACTTGTTTGGTGAAAGGGTTTTACCCGAGCGATATTGCTGTGGAATGGGAGAGTAACGGGCAACCGGAGAACAATTACAAAACGACACCGCCCATGCTTGATAGTGATGGTTCCTTCTTCTTGTACAGCAAGTTGACGGTTGATAAATCCAGGTGGCAGCAAGGAAATGTTTTCTCTTGTTCAGTGATGCATGAGGCGCTCCACAACCATTATACGCAAAAATCACTCTCACTTTCACCGGGGAAATGAAGCTTGAGCAGGGCCT

#### IgG3

CGCATGATGCGTCGACCAAAGGGCCGTCAGTCTTTCCCTTGGCGCCGTGCTCCAGGAGTACCAGCGGCGGCACCGCGGCGTTGGGATGTCTTGTCAAGGATTATTTTCCCGAACCCGTCACCGTAAGCTGGAACAGTGGGGCATTGACGTCTGGCGTTCAT

ACTTTTCCGGCAGTACTTCAGAGTTCCGGCCTTTATTCTTTGTCAAGCGTTGTTACCGTACCATCCAGTAGCCTTGGCACCCAGACCTACACCTGTAATGTTAATCACAAACCAAGTAACACCAAGGTTGATAAGAGGGTTGAGCTTAAAACACCGCTTGGTGACACAACCCATACGTGTCCAAGATGTCCGGAGCCGAAGAGTTGTGATACCCCGCCGCCGTGTCCTCGCTGTCCGGAACCAAAGAGCTGTGATACCCCCCCACCTTGTCCCAGATGTCCTGAACCGAAATCATGTGACACGCCACCACCTTGCCCAAGATGTCCCGCGCCAGAGCTGCTGGGTGGGCCCAGCGTATTTCTTTTTCCACCCAAACCGAAGGATACCCTTATGATAAGCAGGACTCCCGAGGTTACCTGCGTGGTGGTTGACGTAAGTCACGAAGACCCCGAAGTCCAATTCAAATGGTATGTTGATGGGGTCGAAGTACACAACGCGAAGACTAAACCGAGAGAGGAACAGTATAATAGCACATTCCGGGTTGTTTCCGTACTTACAGTACTTCATCAGGACTGGCTTAATGGCAAGGAGTACAAGTGCAAAGTCAGTAACAAGGCACTCCCTGCTCCGATTGAAAAGACAATATCAAAGACGAAAGGTCAACCCAGAGAGCCGCAGGTCTACACACTCCCTCCGTCCAGAGAAGAGATGACGAAAAACCAAGTTTCATTGACGTGCCTCGTTAAAGGATTCTACCCAAGCGACATAGCTGTTGAGTGGGAGAGCAGCGGCCAGCCTGAGAACAATTATAATACTACCCCCCCCATGCTCGACTCTGATGGTAGTTTTTTTCTGTACTCCAAGCTGACGGTAGACAAAAGTAGATGGCAGCAAGGCAACATCTTCAGTTGCTCTGTTATGCACGAGGCGTTGCACAACCGATTCACACAGAAGTCACTGAGCCTGTCTCCGGGTAAATGAAGCTTGAGCAGGGCCT

#### IgA

CGCATGATGCGTCGACTTCTCCAAAAGTGTTTCCCCTCAGTTTGTGTTCCACTCAACCGGATGGTAACGTGGTGATTGCTTGTCTCGTGCAAGGTTTTTTCCCACAGGAACCGCTGAGTGTTACATGGTCAGAGTCAGGCCAAGGTGTAACCGCGCGCAACTTTCCCCCTTCACAGGACGCTAGTGGCGATCTGTATACTACCTCCTCTCAGCTCACTCTTCCCGCCACACAATGCCTCGCTGGGAAATCTGTAACCTGCCACGTTAAACATTACACTA

ATCCATCACAGGACGTTACCGTGCCGTGCCCTGTACCATCCACGCCGCCTACGCCGTCACCGTCAACTCCTCCTACTCCCTCACCCTCTTGTTGTCACCCGCGCCTCTCTCTTCACAGACCGGCCTTGGAGGACCTTCTCCTTGGGTCTGAGGCGAATTTGACTTGCACGCTCACGGGGTTGCGGGACGCTAGTGGGGTTACGTTTACATGGACACCTTCATCAGGGAAGTCTGCCGTTCAGGGCCCCCCAGAGCGCGATTTGTGCGGGTGTTACAGCGTATCTTCTGTGCTGCCTGGGTGCGCTGAGCCCTGGAATCACGGCAAAACGTTTACCTGCACCGCTGCTTACCCAGAGAGCAAAACCCCTCTGACGGCTACATTGTCCAAGTCAGGCAACACATTTCGCCCCGAAGTCCACCTCTTGCCACCTCCATCCGAAGAACTCGCCCTGAACGAACTCGTGACGCTGACGTGCCTTGCACGCGGCTTTTCCCCGAAAGACGTTCTCGTCCGGTGGCTTCAAGGTTCTCAGGAACTCCCACGGGAGAAGTACCTGACCTGGGCTTCACGCCAGGAACCTTCACAAGGGACGACCACTTTCGCAGTCACGTCAATTCTGAGAGTTGCCGCTGAGGACTGGAAGAAGGGAGATACTTTCAGTTGTATGGTAGGTCACGAAGCACTGCCGCTGGCATTTACGCAGAAAACCATCGATCGGCTTGCCGGAAAGCCTACTCATGTTAACGTTTCCGTAGTGATGGCGGAGGTAGATGGCACATGTTACTGAAGCTTGAGCAGGGCCT

### HAI Assays

Sera samples were pre-treated with receptor-destroying enzyme (Denka Seiken, Cat# 370013) followed by hemadsorption, in accordance with WHO recommended protocols (65). HAI titrations were performed in 96-well U-bottom plates (Corning, Mfr# 353077). Sera were initially diluted two-fold and then added to four agglutinating doses of virus, for a final volume of 100 ul/well. Turkey erythrocytes (Lampire, Cat# 7209401) were added to each well (12.5 ul diluted to a 2% v/v). The erythrocytes were gently mixed with sera and virus, then allowed to incubate for one hour at room temperature. Agglutination was read and HAI titers were expressed as the inverse of the highest dilution that inhibited four agglutinating doses of virus. Each HAI assay was performed independently on two different days.

### ‘Headless’ HA ELISAs

‘Headless’ HA ELISAs were performed on 96-well Immulon 4HBX flat-bottom microtiter plates (Thermo Fisher, Cat# 3855) coated with 0.5 ug/well of streptavidin (Sigma, Cat# S4762). Biotinylated ‘headless’ HA protein was diluted in biotinylation buffer containing 1x TBS (Bio Rad, Cat# 170-6435) + 0.005% tween (Bio Rad, Cat# 170-6531) + 0.1% bovine serum albumin (Sigma, Cat# A8022) to 0.25 ug/ml and 50 ul was added per well and incubated on a rocker for one hour at room temperature. Each well was then blocked for an additional one hour at room temperature using biotinylation blocking buffer containing 1x TBS (Bio Rad, Cat# 170-6435) + 0.005% tween (Bio Rad, Cat# 170-6531) + 1% bovine serum albumin (Sigma, Cat# A8022). Each serum sample was serially diluted in biotinylation buffer (starting at 1:100 dilutions for total IgG or 1:50 dilutions for Ab isotype) and added to the ELISA plates and allowed to incubate for one hour at room temperature on a rocker. As a control we added the human CR9114 stalk-specific mAb, starting at 0.03 ug/ml, to verify equal coating of plates and to determine relative serum titers. Next, peroxidase conjugated goat anti-human IgG (Jackson, Cat# 109-036-098), peroxidase conjugated mouse anti-human IgG1 (Southern Biotech, Cat# 9054-05), peroxidase conjugated mouse anti-human IgG2 (Southern Biotech, Cat# 9060-05), peroxidase conjugated mouse anti-human IgG3 (Southern Biotech, Cat# 9210-05), or peroxidase conjugated goat anti-human IgA (Southern Biotech, cat# 2050-05) was incubated for one hour at room temperature on a rocker. Finally, SureBlue TMB Peroxidase Substrate (KPL, Cat# 5120-0077) was added to each well and the reaction was stopped with the addition of 250 mM HCl solution. Plates were extensively washed with PBS (Corning, Ref# 21-031-CM) + 0.1% tween (Bio Rad, Cat# 170-6531) between each step using a BioTek microplate washer 405 LS. Relative titers were determined using a consistent concentration of the CR9114 mAb for each plate and reported as the corresponding inverse of the serum dilution that generated the equivalent OD. Each type of ELISA (total IgG, IgG1, IgG2, IgG3, and IgA) was performed twice.

### Chimeric (c6/H1) HA ELISAs

Chimeric HA ELISAs were performed on 96-well Immulon 4HBX flat-bottom microtiter plates (Thermo Fisher, Cat# 3855). HA proteins were diluted in PBS (Corning, Ref# 21-031-CM) to 2 ug/ml and coated at 50ul per well overnight at 4C. Plates were blocked using an ELISA buffer containing 3% goat serum (Gibco, Cat# 16210-064) + 0.5% milk (dot scientific inc., Cat# DSM17200-1000) + 0.1% tween (Bio Rad, Cat# 170-6531) in PBS (Corning, Ref# 21-031-CM) 1x for two hours at room temperature. Each serum sample was serially diluted in the ELISA buffer (starting at 1:100 dilutions) and added to the ELISA plates and allowed to incubate for two hours at room temperature. As a control we added the human CR9114 stalk-specific mAb, starting at 0.03 ug/ml, to verify equal coating of plates and to determine relative serum titers. Next, peroxidase conjugated goat anti-human IgG (Jackson, Cat# 109-036-098) was incubated for one hour at room temperature. Finally, SureBlue TMB Peroxidase Substrate (KPL, Cat# 5120-0077) was added to each well and the reaction was stopped with the addition of 250 mM HCl solution. Plates were extensively washed with PBS (Corning, Ref# 21-031-CM) + 0.1% tween (Bio Rad, Cat# 170-6531) between each step using a BioTek microplate washer 405 LS. Relative titers were determined using a consistent concentration of the CR9114 mAb for each plate and reported as the corresponding inverse of the serum dilution that generated the equivalent OD. Each ELISA was performed twice.

### Competition ELISAs

Competition ELISAs were performed on 96-well Immulon 4HBX flat-bottom microtiter plates (Thermo Fisher, Cat# 3855). HA proteins were diluted in DPBS 1x (Corning, Ref# 21-031-CM) to 2 ug/ml and coated at 50ul per well overnight at 4C. Plates were blocked using the biotinylation blocking buffer, described earlier, for two hours at room temperature. Each serum sample was serially diluted in biotinylation buffer (starting at 1:10 dilution) and added to the ELISA plates and allowed to incubate for one hour at room temperature before adding the human 70-1F02 mAb (specific for the conformationally dependent HA stalk epitope 1 (66) that had been biotinylated using the Invitrogen SiteClick Biotin Antibody Labeling Kit (Thermo Fisher, Cat# S20033) at a constant concentration of 0.03 ug/ml and incubated at room temperature for an additional hour. As a control we added the human CR9114 stalk-specific mAb, starting at 0.03 ug/ml, to verify equal coating of plates and to determine relative serum titers. Next, peroxidase conjugated streptavidin (BD Pharmingen Cat#554066) was incubated for one hour at room temperature. Finally, SureBlue TMB Peroxidase Substrate (KPL, Cat# 5120-0077) was added to each well and the reaction was stopped with the addition of 250 mM HCl solution. Plates were extensively washed with PBS (Corning, Ref# 21-031-CM) + 0.1% tween (Bio Rad, Cat# 170-6531) between each step, with the exception of the addition of the biotinylated 70-1F02, using a BioTek microplate washer 405 LS. Relative titers were determined using the un-competed control lane OD (biotinylated 70-1F02 binding in the absence of sera) and setting that OD as 100%. Each serum sample was then assessed by non-linear regression using GraphPad Prism. Titers are reported as the inverse of the highest serum dilution that inhibited binding of the biotinylated 70-1F02 to 30% of the un-competed binding. Each competition ELISA was performed independently on two different days.

### *In Vitro* Neutralization Assays

Plasmids encoding pH1N1 viruses possessing genes encoding enhanced green fluorescent protein (eGFP) in place of most of the PB1 gene segment were provided by Jesse Bloom at The Fred Hutchinson Cancer Research Center. The eGFP segment retained the noncoding and 80 terminal coding nucleotides, allowing this segment to be efficiently and stably packaged into the virions. Detailed protocols for the reverse genetics, expression, and *in vitro* neutralization assays using the recombinant viruses have been published elsewhere (59, 67). In brief, serum was pre-treated with receptor-destroying enzyme (Denka Seiken, Cat# 370013) and then serially diluted in neutralization assay media (Medium 199 (Gibco, Cat# 11150-059) supplemented with 0.01% heat inactivated FBS (Sigma, Cat# F0926-100) + 0.3% bovine serum albumin (Sigma, Cat# A8022) + 100 U penicillin/100 ug streptomycin/ml (Corning, Cat# 30-002-Cl) + 100 ug of calcium chloride/ml (Sigma, Cat# S7653) + 25 mM HEPES (Corning, Cat# 25-060-Cl)), beginning at 1:80 dilution. PB1flank-eGFP viruses were then added to the sera dilutions and were incubated at 37C for one hour to allow for neutralization. As a control, the human CR9114 HA stalk-specific mAb was added to ensure equal infectivity and neutralization across all plates. Viruses and sera were then transferred to 96-well flat-bottom tissue culture plates containing 80,000 cells per well of MDCK-SIAT1-TMPRSS2 cells constitutively expressing PB1 under a CMV-promoter. Plates were incubated at 37C for 30 hours post-infection. Mean fluorescent intensity of samples was read using an Envision plate reader (monochromator, top read, excitation filter at 485 nm, emission filter at 530 nm). Neutralization titers were reported as the inverse of the highest dilution that decreased mean fluorescence by 90%, relative to infected control wells in the absence of antibodies. Each neutralization assay was performed independently on two different days.

### ADCC activity Assays

293T cells plated at 3.5e4 cells per well in a 96-well flat-bottom tissue culture plate (Corning, Cat# 353072) 24 hours before transfection. 293T cells were then transfected using 20 ul OptiMEM (Gibco, Cat# 31985-070) + 1 ul Lipofectamine 2000 (Invitrogen, Cat# 11668-019) + 500 ng plasmids encoding the HA gene from A/California/07/09 per well and incubated at 37C for approximately 30 hours before performing the ADCC assay. Approximately 12 hours before performing the ADCC assay, frozen PBMCs from four separate donors (obtained through the University of Pennsylvania Human Immunology Core) were thawed at 37C and then washed 3x using 15 mls of warmed complete RPMI media (Corning, Cat# 10-040-CM) (supplemented with 10% heat-inactivated FBS (Sigma, Cat# F0926-100) + 1% Penn-Strep (Corning, Cat# 30-002-Cl)). Each aliquot of PBMCs was then transferred to a 50 ml conical and rested overnight in 23 mls of complete RPMI media at a 5 degree angle with the cap loosened to allow for gas exchange. On the day of the assay, sera was diluted in DMEM (Corning, Cat# 10-013-CM) supplemented with 10% FBS (Sigma, Cat# F0926-100) at a 1:10 dilution. As a control for this assay, the human CR9114 HA stalk-specific mAb was included at a concentration of 5 ug/ml to ensure efficient activation of ADCC. Transfected 293T cells were loosened by pipetting and transferred to a 96-well U-bottom plate (Corning, Mfr# 353077), spun down for 1 minute at 1200 RPM, and the media was flicked out. The sera/mAb dilutions were transferred to the plates containing the transfected 293T cells and were mixed with the transfected cells by gentle pipetting and incubated at 37C for two hours. PBMC aliquots were combined, spun down, counted, and a master mix of 2e7 cells/ml was set up using complete RPMI media. Aliquots of the PBMC master mix were set up for the live/dead and unstained control wells. PE-conjugated mouse anti-human CD107a (BioLegend, Cat# 328608) was added at a 1:50 dilution. Brefeldin A (Sigma, Cat# B7651) was added to 10 ug/ml. Monensin (BD BioSciences, Cat# 51-2092KZ) was added to 5 ul per 1 ml of PBMC master mix concentration. An aliquot of 200 ul was made and PMA (Sigma, Cat# P1585) was added to a 5ug/ml concentration and ionomycin (Sigma, Cat# I9657) was added to a 1 ug/ml concentration. Serum/cell suspensions were spun down at 1200 RPM for 1 minutes and media was flicked out. The PBMC master mix and the aliquots for the various controls were plated at 50 ul per well and mixed gently by pipetting, followed by incubation at 37C for four hours. Cells were then stained in the following manner. Live/Dead fixable near-IR stain (Thermo, Cat# <u>L34976)</u> was diluted 1:50 in DPBS (Corning, Ref# 21-031-CM) + 1% bovine serum albumin (Sigma, Cat# A8022) for 30 minutes in the dark at 4C. Human FcR blocking reagent (Miltenyi Biotec, Cat# 130-059-901) was diluted 1:25 in DPBS (Corning, Ref# 21-031-CM) + 1% bovine serum albumin (Sigma, Cat# A8022) and incubated in the dark for 10 minutes at 4C. AlexaFluor 647-conjugated mouse anti-human-CD3 (BioLegend, Cat# 344826) and BV421-conjugated mouse anti-human CD56 (BioLegend, Cat# 318328) were diluted 1:200 in DPBS (Corning, Ref# 21-031-CM) + 1% bovine serum albumin (Sigma, Cat# A8022) and incubated in the dark for 30 minutes at room temperature. Cells were then fixed using 10% paraformaldehyde (Electron Microscopy Sciences, Cat# 15714-S) diluted in milliQ water for 6 minutes at room temperature. Cells were extensively washed with DPBS (Corning, Ref# 21-031-CM) + 1% bovine serum albumin (Sigma, Cat# A8022) between each step. Cells were stored overnight at 4C in 100 ul/well of DPBS (Corning, Ref# 21-031-CM) + 1% bovine serum albumin (Sigma, Cat# A8022). Flow cytometry was performed using (LSRII, BD Biosciences, San Diego, CA). Compensation controls were set up using anti-mouse Ig κ beads (BD BioSciences, Cat# 552843) and run for each antibody for every experiment and voltages were adjusted accordingly. All data were analyzed in FlowJo (Ashland, OR), by gating on single cells that were CD3^-^/CD56^+^/CD107a^+^ in control wells that did not contain serum/mAb to adjust for basal levels of CD107a expression. These gates were then applied to each serum sample and ADCC activity was expressed as the fold-change over background. Each ADCC assay was performed independently on three different days and the same four PBMC donors were pooled and used for each replicate.

### Murine Experiments

All passive transfer experiments were performed in humanized FcR mice (hFcgR (1, 2a, 2b, 3a, 3b)tg^+^/mFcgR alpha chain (1, 2b, 3, 4)^-/-^) that were provided by Jeff Ravetch at The Rockefeller University (61). Sera were pooled into three groups: uninfected-HAI^high^ (>40 HAI titer), uninfected-HAI^low^ (≤ 40 HAI titer), and infected-HAI^low^ (≤40 HAI titer), and then heat-treated for 30 minutes at 55C. Sera or sterile PBS was then transferred into mice by intraperitoneal injection. Four hours post transfer, mice were bled by sub-mandibular puncture and then anesthetized using isofluorane and challenged intranasally using 50 ul of sterile PBS containing a sublethal dose (9e4 TCID50 units) of A/California/07/09. ELISAs were run on the sera collected from each animal to verify the passive transfer was successful. Mice were weighed on the day on infection and then daily x15 days post infection. Weight loss was reported as percent weight loss relative to the starting weight of each mouse. For the passive transfer normalized by volume, two independent experiments were performed using a mix of male and female humanized FcR mice for a total of 6 mice per group per experiment. For the passive transfer normalized by HA antibody titer, a single experiment was performed using a mix of male and female humanized FcR mice for a total of 6 mice per group, since we had limited amounts of sera available for this study. One-way ANOVA was performed for each day post-infection between groups using GraphPad Prism software.

### Statistical Analysis

Fisher’s exact tests and one-way ANOVAs were completed using GraphPad Software (2018). Both unadjusted and adjusted logistic regression analyses were performed using R Studio (Version 1.0.153).

## Acknowledgments

This work was supported by the National Institute of Allergy and Infectious Diseases (1R01AI113047, SEH; 1R01AI108686, SEH; 1R01AI097150, ASM; CEIRS HHSN272201400005C, SEH and ASM) and Center for Disease Control (U01IP000474, ASM). Scott E. Hensley holds an Investigators in the Pathogenesis of Infectious Disease Awards from the Burroughs Wellcome Fund. We thank Florian Krammer (Mt. Sinai), Jesse Bloom (Fred Hutchinson Cancer Center), Patrick Wilson (University of Chicago), Barney Graham (Vaccine Research Center, NIH) and Adrian McDermott (Vaccine Research Center, NIH) for providing plasmids for this study.

---

**Supplemental Table 1 – Demographic characteristics of subjects enrolled in hospital-based human cohort study.** The second and third column list the number and percentage of each demographic characteristic within the infected and uninfected groups, respectively.

**Supplemental Table 2 – One-way ANOVA results for passive transfer normalized by volume (Fig. 4B).** One-way ANOVA with Tukey’s correction for multiple comparisons performed for each dpi. The p-value for each comparison is reported, calculated based on the %-weight lost compared to baseline for each group, each day.

**Supplemental Table 3 – One-way ANOVA results for passive transfer normalized by HA titer (Fig. 4D).** One-way ANOVA with Tukey’s correction for multiple comparisons performed for each dpi. The p-value for each comparison is reported, calculated based on the %-weight lost compared to baseline for each group, each day.

**Supplemental Figure 1 - HAI titers of >40 are significantly associated with decreased risk of influenza infection. *S1A:*** HA head-specific Ab titers against the circulating H1N1 strain were determined by HAI. Uninfected individuals had significantly increased HAI titers compared to infected individuals (p = 0.0008, two-tailed Fisher’s Exact Test based on the geometric mean titer of two independent experiments).

**Supplemental Figure 2 - Isotype swapping of HA stalk mAb CR9114.** The constant region IgG1, IgG2, IgG3, or IgA of mAb CR9114 were cloned into the vector expressing the heavy chain of CR9114. The variable region (heavy and light chains) were maintained between each clone to ensure equivalent antigen recognition and binding. Each isotype was sequence-confirmed, expressed, and purified for use in ELISAs.

